# Drawing the experience dynamics of meditation

**DOI:** 10.1101/2022.03.04.482237

**Authors:** Barbara Jachs, Manuel Camino Garcia, Andrés Canales-Johnson, Tristan A. Bekinschtein

**Affiliations:** Cambridge Consciousness and Cognition Lab, Department of Psychology, University of Cambridge, UK; Department of Developmental and Educational Psychology, Faculty of Psychology. University of Valencia, Spain; Vicerrectoría de Investigación y Posgrado, Universidad Católica del Maule, Talca, Chile

## Abstract

Subjective experiences are hard to capture quantitatively without losing depth and nuance, and subjective report analyses are time-consuming, their interpretation contested. We describe Temporal Experience Tracing, a method that captures relevant aspects of the unified conscious experience over a continuous period of time. The continuous multidimensional description of an experience allows us to computationally reconstruct common experience states. Applied to data from 852 meditations – from novice (n=20) and an experienced (n=12) meditators practising Breathing, Loving-Kindness and Open-Monitoring meditation – we reconstructed four recurring experience states with an average duration of 6:46 min (SD = 5:50 min) and their transition dynamics. Three of the experience states assimilated the three meditation styles practiced, and a fourth experience state represented a common low-motivational, ‘off-task’ state for both groups. We found that participants in both groups spent more time in the task-related experience state during Loving Kindness meditation than other meditation styles and were less likely to transition into an ‘off-task’ experience state during Loving Kindness meditation than during Breathing meditation. We demonstrate that drawing the dynamics of experience enables the quantitative analysis of subjective experiences, transforming the time dimension of the stream of consciousness from narrative to measurable.

## Introduction

In 1996, Francisco Varela proposed the research program of neurophenomenology as a methodological remedy for the hard problem of consciousness^1^. At the heart of this program stands the necessity for the “[…] re-discovery of the primacy of human experience and its direct, lived quality […] “^1^, as only a balanced account of the external and the experiential perspectives would allow progress on the questions of why and how phenomenal consciousness exists. By studying the phenomena of experiences with the same rigour as the neurophysiological and behavioural phenomena, we can characterise these phenomena, their dependencies, dissociations and causal structures. This requires the development of rigorous methods that enable the exploration and analysis of lived first-person experience^2,3^.

Phenomenological methods that have been developed for cognitive research, can be broadly categorised into the traditional phenomenological interview techniques (for a discussion of these methods, see^4^) and methods of front-loading phenomenology into the experimental design^4–7^, such as the experience sampling method^8–10^.

In order for phenomenological data to be integrated into neuroscience research, a primary aim needs to be the capture of non-idiosyncratic (i.e. intersubjectively valid) aspects of experience that can guide the analysis of neurophysiological data^3,9,11–14^. In a landmark paper, Lutz et al. (2002)^11^ trained participants to describe their pre-trial state during a behavioural task. These descriptions were clustered into phenomenologically distinct levels of “preparedness” and explained a significant portion of observed variance in task performance and EEG brain activity.

Methods with a wider application potential may require the “front-loading” of phenomenology into the experimental design^5^. For example, in two recent studies, Canales-Johnson *et al*. ^15,16^ dissociated two phenomenological variants of the same stimulus during an auditory and a visual bistable perception task using frontoparietal information dynamics. In this experimental design the participants were asked to discriminate between two pre-defined, phenomenologically distinct experience phenotypes known to be evoked by the ambiguous stimulus. By front-loading the phenomenology into the experimental design, the researchers were able to study the neural correlates associated with the experience of each phenomenological class. This demonstrates how obtaining phenomenologically distinct, intersubjectively reliable classes of experience can allow for phenomenology to inform cognitive neuroscience research.

A framework conceptualising the contents of thought is the stream of consciousness, formalized by William James ^17^, which can be understood as the continuous flow of dynamic integrated experiences. James observed that “personal consciousness thought is always changing” and that it is “sensibly continuous” and as such comparable to a river or stream^17^. Similar analogies exist in early Buddhist scriptures, describing sensory experiences and mental events, such as feelings, perceptions and volitions, to flow in a constantly changing “mind stream”^18^. It remains a challenge to reliably capture the stream of consciousness, which would enable the study of the dynamics of continuous experiences, such as altered states of consciousness, flow states^19^, or the affective states associated with mental disorders^20^.

The history of experimental psychology^21^ and cognitive neuroscience^22^ shows that it is customary to capture the dynamics of neurophysiological measures such as EEG and fMRI and/or behavioural responses like reaction times and eye movements, but not the dynamics of experience^23^. We have been able to find only one project that used a retrospective timeline to capture perceived meditation depth (Müller, 1997^24^). By relating the meditation depth diagram to a meditation experience questionnaire, the study found the definition of meditation depth to differ between participants. Outside of cognitive neuroscience, a similar technique named “timelining” is frequently used in product and policy design to understand a consumer’s perceived satisfaction with a product over time and to understand a typical user’s journey^25^. This is also frequently used as a method in life history research^4^. Inside cognitive sciences, autobiographical memory research takes the passing of time and the experiences and events in sequence as a primary tool to study its mechanisms^26^.

In this study we introduce the temporal experience tracing method (TET) by applying it to the investigation of the dynamics of phenomenological features of three commonly practiced styles of meditation – mindfulness of breathing (B), loving-kindness meditation (LK), and open monitoring (OM) – to evoke different mental states, despite the absence of behavioural differences.

We defined relevant aspects of meditations’ experiences in conversation with expert meditators and, after each meditation, participants graphically represented the changes in intensity of each dimension of experience (**Figure 1**), creating a twelve-dimensional description of their experience state in time. Half of the phenomenological dimensions chosen for this study were taken from the recently proposed phenomenological matrix of mindfulness meditation, which draws upon the psychotherapeutic literature, traditional Buddhist accounts, contemporary meditation manuals and scholarly analyses of traditional mindfulness meditation^27^. This matrix was an attempt to answer the question: *“… when one is engaged in a formal mindfulness practice, what observable, instructible, and manipulable features of experience are most relevant to training in mindfulness?*” ^27^. Some aspects of experience were not included in the original matrix, as they represent shared contextual features of mindfulness meditation styles, and as such would be less significant in distinguishing between meditation styles^28^.

**Figure 1:**
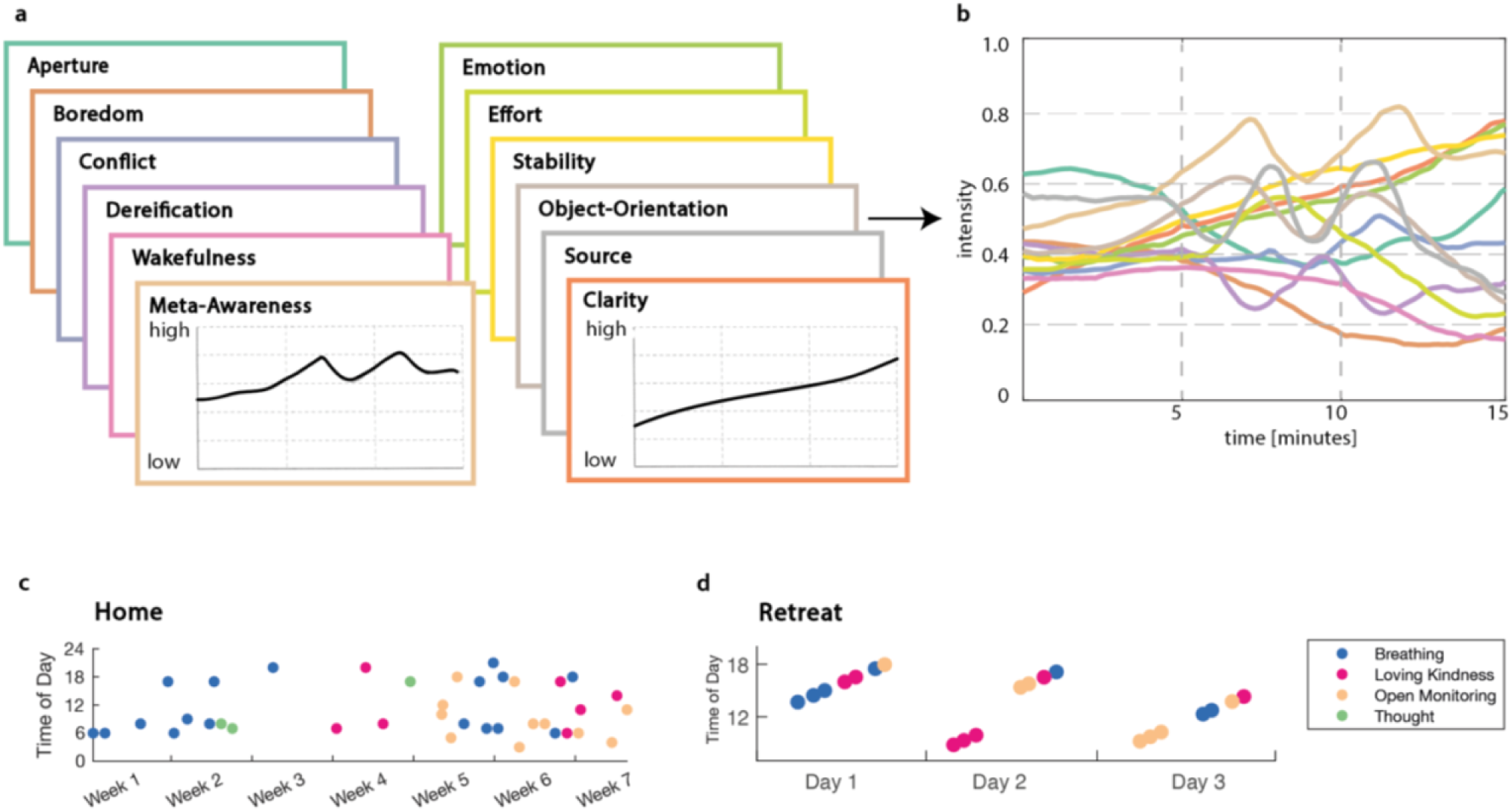
The experimental design using temporal experience tracing (TET). After each session, participants graphed their experiences along twelve dimensions in the temporal experience traces (TET). **a** – example TETs from one retreat meditation session. **b** – Once digitised the data generate a multidimensional experience profile in time. The data can now be analysed using any conventional timeseries analysis. Two groups of meditators completed the temporal experience traces after each meditation session. **c** – An example of adherence by a participant of the home group (n=12) who recorded 38 meditation sessions. **d** – Participants in the retreat group (n=20) meditated 21 times (7 sessions meditation style) over three days.

In this project, we were interested in capturing all potentially salient aspects of experience during meditation, and we therefore included the additional dimensions “Emotion”, “Conflict”, “Wakefulness”, “Boredom” and “Source”. These are closer to psychological constructs used in basic and clinical cognitive neuroscience and hence transferable. While not necessarily primary targets for the meditation practice, these dimensions may show substantial variance across sessions and affect other cognitive experience dimensions. For example, fluctuations in daytime wakefulness, while part of normal brain functioning, are known to affect cognitive processing ^29–31^ and phenomenology ^32,33^. More detail on each dimension can be found in the methods section.

Two meditation groups used the temporal experience tracing method (TET) to report on their experiences after each meditation they either practiced at home (in the case of the Home Group) or at a meditation retreat (the Retreat Group). We collected a minimum of 21 meditation sessions per participant, giving this study has a powerful repeated-measures between-group design. We identify recurring clusters of dimension intensity combinations, and examine their dynamics across individuals, meditation styles and meditation groups. This analysis is rooted in the concept that the experience dimensions, while they can be articulated separately, are likely mutually affecting each other, and therefore cannot be thought of as independent from each other^34^. Additionally, many theories of consciousness are agreeing on the indivisibility, irreducibility of conscious experience^35–37^. We therefore computationally devise combinations of dimension intensities that are common across participants to approximate common intersubjective classes of experience during mindfulness meditation practice. Finally, we use Markov Chains to study the dynamics of transitions between experience states. We show that this approach can capture (i) recurring metastable experience states, (ii) the temporal dynamics of these experience states in three meditation styles, and (iii) experiential commonalities and differences between meditation styles across participants.

## Methods

### Participant Groups

Two datasets were collected as part of a neurophenomenology study, including low-density portable EEG recordings and behavioural data^38^. These are referred to as Retreat dataset and Home dataset. The Retreat dataset was collected over two mindfulness retreats with ten novice meditators participating in each retreat. The retreats took place over three days near Valencia, Spain, where the meditators were led through seven meditations of each style - Breathing Meditation, Loving Kindness Meditation and Open Monitoring Meditation. Participants had been practicing mindfulness meditation for less than one year and did not have a regular practice. Each meditation was 15 minutes long and a gong was played at the beginning-, five-, ten- and 15-minute marks.

The Home dataset consists of twelve participants, who recorded a minimum of 35 meditations at home. The participants were recruited from the Jamyang Buddhist Centre and Buddhist Society in London and were admitted to the study if they had an established regular meditation practice and were familiar with four meditation styles (Samatha of Breathing, Samatha of Thought (Focused Attention styles), and Samatha without an Object (Open Monitoring), as well as Meta Bhavana (Loving Kindness) meditation). Participants had an estimated mean 4.2 years of meditation experience (STD=3.36y).

Meditation practices at home were guided by audio recordings, which we made available on the online platforms SoundCloud and Insight Timer. These guides were recordings of a script provided by the meditation teacher. Each meditation was 20 minutes long and started with a five-minute body scan. A gong sounded every five minutes and was followed by brief guiding (∼14 seconds) for one of the four meditation styles. An additional recording contained the gong sounds in 5-minute intervals only, if participants preferred to meditate without additional guidance. No restrictions were made on the time of day or sequence of the meditation styles to practice, minimising any changes to their meditation practice the experiment caused, but participants were asked to practice all meditation styles across the course of the experiment.

### The Dimensions

To ensure that the dimensions participants use to represent their experiences are meaningful to the researchers and participants, the experience dimensions were developed in collaboration with the meditation teachers who were instructing the respective classes participating in the study. The dimensions used were based on the theoretical framework of the phenomenological matrix for meditation experiences^27^. In this framework, the three primary dimensions are hypothesised to be orthogonal to each other: object orientation, Dereification and meta-awareness. Further, four secondary dimensions were proposed to describe highly relevant features of experience that are affected by mindfulness practices: Aperture, clarity, stability, and effort ^27^.

This framework was further adapted in collaboration with Dr Elizabeth English, who teaches mindfulness for students at the University of Cambridge and Roy Sutherwood, who led the meditation groups at the Buddhist Society. In this process, the five additional dimensions of wakefulness, stability, source, conflict and emotion were added, to capture typical challenges experienced by novice meditators. The results were the following twelve dimensions: Meta-Awareness, Boredom, Clarity, Conflict, Dereification, Effort, Emotion, Object Orientation, Aperture, Source, Stability and Wakefulness. A Spanish version was developed for the Mindfulness Retreat participants, who were native Spanish-speakers.

#### Attention-related Dimensions

##### 1. Meta-Awareness

Meta-Awareness refers to the monitoring of mental activities and of contents of consciousness^3,27^. Noticing mind-wandering during meditation is an example of meta-awareness and is therefore part of most meditative practices. It is thought to be particularly relevant in Open Monitoring meditation, where awareness is maintained even if the meditator engages in the thoughts and experiences that arise.

##### 2. Object Orientation

Object orientation refers to the cognitive concept of endogenous orienting or top-down attention. During mediation it reflects the degree to which the mental state is oriented toward a specific object or class of objects, such as a physical sensation (e.g., the breath), a visualisation or a mantra. It has therefore been stipulated that object-orientation is high during focused attention style meditations^39^, such as Breathing or Loving Kindness meditation. Low object-orientation might be found when the meditator gets distracted during focused attention meditation. Additionally, Open Monitoring meditation might encourage low Object Orientation, as they deliberately reduce the engagement with any particular object of attention^27^.

##### 3. Aperture

Aperture refers to the broadness of the scope of attention, as described in the classical optical analogy of the “spotlight of attention”. Low Aperture would reflect a narrow spotlight of attention that focuses on a specific object, for example the tip of the nostril during a focused attention meditation on breathing. High Aperture would refer to a wide attentional field, such as practiced in OM meditation styles^27^.

##### 4. Stability

This dimension reflects the continuity of a state over time despite internal or external perturbations. During meditation, high stability can be achieved through sustained attention on the object or field of attention^40^. However, rumination can also be a stable state over time. Low stability reflects the experience of fluctuating mental states^27^. Neurophysiologically, meditation as a state and expertise has been linked to increased EEG microstate stability of resting-state networks^41^, but it remains unclear to what degree this measure reflects the subjective experience of stability.

#### State-related

##### 5. Wakefulness

Drowsiness is a common hindrance in novice meditators. During the process of falling asleep a progressive impairment of each step of cognitive processing is likely to reduce the meditators ability of task-retention^42^. High Wakefulness reflects the meditators subjective experience of wakeful arousal. The lowest score on this scale corresponds to having fallen asleep.

#### Motivation-related dimensions

##### 6. Boredom

Another frequent hindrance, boredom, is common in meditators^43^. Phenomenologically, it is an unpleasant and distressing experience, frequently comprising of feelings of restlessness and lethargy^44^. It is thought to arise of a failure of engagement of executive control networks when faced with a monotonous, uninteresting task^45^.

##### 7. Effort

Effort refers to the subjective experience of sustaining a high cognitive workload despite difficulties, and may therefore be related to boredom^44^. The ability to sustain effortful states has been found to be moderated by task difficulty, trait Conscientiousness^46^ and competence belief^47^. Physiologically, effort is thought to reflect the subjectively experienced negative valence tracking the cost of allocating working memory to a specific task^48^. In meditation, early stages of meditation are thought to require more effort than later stages, when the meditator can modulate how much effort, or cognitive resource, to dedicate to the practice^27,49^.

#### Content-related dimensions

##### 8. Clarity

Clarity is the subjective experience of vividness and unambiguity of the stream or contents of consciousness. In the proposed phenomenological matrix of meditation, Lutz et al compare this aspect of internal experience to the ability of attention to change the contrast appearance of visual gratings^27^. In this way, high attention states, such as achieved during focused attention meditation would be associated with more clarity than the contents of consciousness during mind wandering. Additionally, other less salient elements of experience, such as the underlying affect, may turn into distinct features when high clarity and meta-awareness combine.

##### 9. Conflict

The subjective experience dimension of conflict captures the extent to which meditators feel conflicted about the content of consciousness. In novice meditators this can be high, when participants judge their own thoughts and experiences, and their ability to adhere to the task^50,51^. In mindfulness meditation the feeling of conflict, or lack thereof, can dissociated from other emotions. For example, a meditator might feel anger or sadness while being accepting of those feelings, experiencing little conflict. Alternatively, a meditator might experience positive emotions, but by judging them, create internal conflict.

##### 10. Dereification

The dimension of Dereification refers to the experience of the cognitive understanding that the thoughts, feelings and perceptions are not the same as reality, but rather a mental process. This process is also known as phenomenological reduction^1^. In the previously proposed phenomenological matrix of meditation, this is elucidated by the example whereby a meditator has the thought “I am a failure” – this thought can be experienced as an accurate depiction of reality, in which case the experience is highly reified, or as merely a thought without representational integrity^27^.

##### 11. Emotion

Here participants are asked to report on the intensity of the emotions they are experiencing, regardless of their valence. This includes the perceived affective tone of their mental state, affective mental events and actively generated emotional content.

##### 12. Source

Different meditation styles have different attitudes towards the meditator playing an active or passive role in creating or manipulating the contents of consciousness. A high intensity of source refers to the meditator taking an active role in generating the content, whereas a low intensity of source reflects a passive stance, whereby mental content is allowed to arise and pass without any active engagement with it^52^.

### Application

Participants were given detailed instructions on how to use the experience traces. For each dimension, participants were given written descriptions and examples. In both experimental setups, participants had the opportunity to discuss the dimensions with the experimenters and teachers for clarification.

After each meditation, participants were given empty graphs with horizontal and vertical grid lines. The vertical grid lines were placed to the five-minute mark, to match the gongs in the audio guides. Two participants in the Home meditation group completed the graphs digitally (OneNote, Microsoft), while the rest of the participants used a pen-and-paper version of the task, drawing into a printed booklet. The retreat dataset contains 399 meditation sessions collected from 19 participants over three days. Included in the Home dataset are 467 meditation sessions collected from 12 participants over a mean period of 102.69 days (Std=62.98 days).

### Pre-processing

The image files of the pen-and-paper and digital experience traces were vectorised using a custom MATLAB script. The accuracy of the digitisation was manually checked, and the scanned images were edited where necessary to increase faithfulness of digitisation. Sessions with missing dimensions were removed.

### Dimension Intensities

We conducted a 2-way analysis of variance (ANOVA) to test the effects of the factors Group (Home or Retreat) and Meditation Style (Breathing, Loving Kindness, Open Monitoring), on the mean intensity of each subjective experience dimension. We corrected for multiple comparison using the Benjamini-Hochberg critical value^53^ assuming a false discovery rate of 10%. To compare the sizes of effects in the study we computed the partial eta squared (η_p_^2^), which is the ratio of variance associated with an effect to the sum of the variance associated with an effect and its associated error variance^54^.

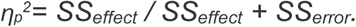

Partial eta squared improves the comparability of effect sizes between studies when more than one variable is being manipulated^55^.

### Cluster Analysis of the experience traces

Clustering algorithms partition existing data into similarity or proximity-based groupings. In this case, the clustering algorithm partitioned the time points into clusters based on their similarity in all 12 experience dimensions. This is not a data reduction technique, instead the clustering allows us to detect natural groupings in the data, without any *a priori* knowledge of group labels.

In this case, the cluster analysis allows us to detect clusters of similar reported experiences across all participants and meditation sessions, and to compare their frequency between participants, meditation styles and between groups of participants.

The time series data were clustered (k-means) using squared Euclidean distance as distance metric. The number of clusters was selected visually (‘elbow method’) from the scree plot of the sum of squared errors that result from clustering the data into an increasing number of clusters (k). The optimal number of clusters for the data was selected as the point after which the inertia decreases visibly and the return for adding a further cluster becomes small. We performed an additional exploratory analysis with clustering using a smaller and larger number of clusters (k).

The exact clustering solutions resulting from k-means clustering are dependent on the initial position of the cluster centroids. We therefore replicated the clustering (n=1000) with randomly chose n initiation points. The clustering with the lowest resulting sum of distances was selected for further analyses.

The clustering solution for *k = 4* is visualised in *Supplementary Figure S3*, where each of the datapoints of the timeseries (101 time points per meditation session) is coloured by its cluster identity. The k-means clustering solution is visualised in the first three principal component space. The clustered data was averaged for each participant to allow visualisation of the distances between participants within each cluster using Multidimensional scaling (MDS)^56^. This visualisation showed a clear separation between the clusters along the first three MDS dimensions.

To understand the properties of each cluster, we conducted a separate two-way ANOVA on the intensity of each dimension, to assess the factors ‘group’ and ‘cluster identity’ as sources of variance, correcting for multiple comparison using the Benjamini-Hochberg critical value and calculating the partial effect sizes.

Once the clusters were computed, we assessed whether the relative frequency of each cluster within each of the meditation styles was different between the two Groups of Participants practising at Home or on the Retreat. The proportion of time spent in each cluster was computed for each participant and square root transformed to correct for the positive skew and unequal variances. The corrected datasets passed Mauchly’s test for sphericity and followed a normal distribution.

To understand the effects of cluster identity, meditation style and meditation group on the relative amount of time spent within each experience state, we set up a repeated-measures ANOVA (Within: clusters, meditation style; Between: groups). Posthoc tests were corrected for multiple comparison using the Benjamini-Hochberg critical value.

Finally, we computed the transition matrices for each meditation style and participant group. The transition matrix is defined by *W*_*i,j*_ in which *i* is the current state, and *j* the state transitioned into. From these transition matrices we built discrete-time, finite-state, time-homogeneous Markov Chains. In a Markov chain the current state contains all the information necessary to predict the likelihood of the next step in the chain^57^.

Using these Markov Chains, we then examined whether the likelihood of transition between clusters depended on meditation style and group. Furthermore, these simple Markov Models allows us to then run simulations from even starting distributions to understand the tendencies of the system at their stability points.

## Results

### The correlations between experience dimensions show internal consistency

The correlation matrices computed between experience dimensions for each session (**Figure 2**) showed high similarity between meditation styles (*Mantel-z test; Retreat: r (B*|*LK) = 0*.*762; r (B*|*OM) = 0*.*649; r (LK*|*OM) = 0*.*672; Home: r (B*|*LK) = 0*.*863; r (B*|*OM) = 0*.*794; r (LK*|*OM) = 0*.*610, p<0*.*05)*. On the individual participant level however, 12 of the 19 retreat group participants did not have a reliable similarity in at least one combination of meditation styles. In the home group, this was the case in 5 of the 12 participants. Therefore, while on the group-level the relationship between dimensions stayed consistent, on an individual participant level the relationships between dimensions changed depending on the context of the meditation style being practiced.

**Figure 2.**
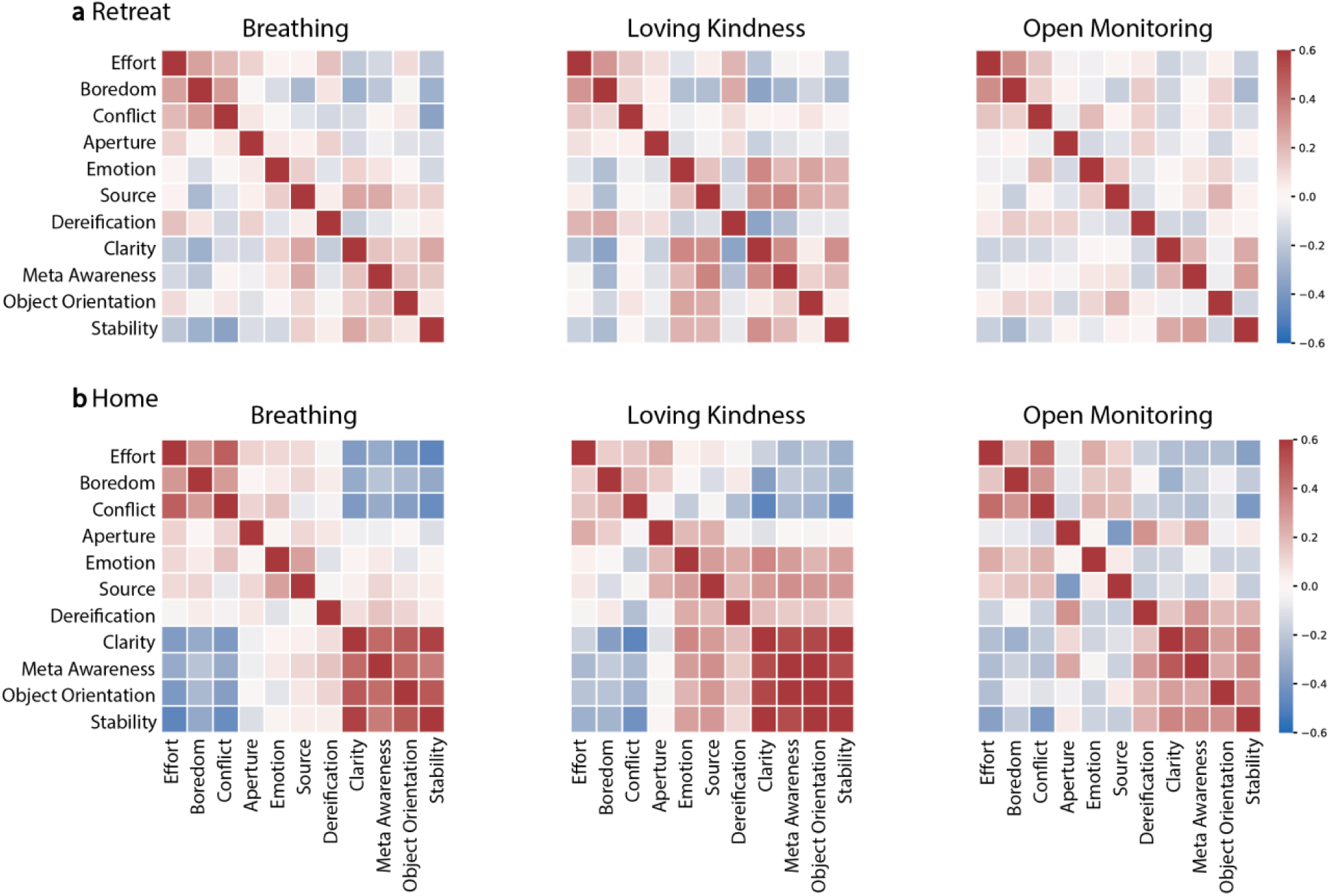
Correlation matrices suggest the existence of clusters in the dimensions which may describe similar experiences. averaged across participant **Top panel** shows the data from retreat group participants, **bottom panel** shows data from home group. Stronger correlations in the Home group, as well as higher similarity of correlation matrices across meditation styles suggest a higher consistency of experience reporting, and less variance introduced by the individual meditation styles context.

Interestingly, the correlation matrices revealed natural groupings within the data. The dimensions (i) Effort, Boredom and Conflict were positively correlated across all conditions, as were (ii) Clarity, Meta-Awareness and Stability. Such grouping within correlation matrices is indicative of natural clusters within the data^58^. This was corroborated by a principal component analysis, which revealed that the dimensions were not orthogonal, with some dimensions loading highly on multiple components, indicating nonmonotonic or nonlinear relationships between experience dimensions. Taken together, this suggests an inherent structure to the data, whereby certain combinations of dimension intensities are more common than others. This is in line with contemporary theories of consciousness that agree on the integrated nature of conscious experience^35,36^. Under such a framework, each experience dimension describes a different aspect of the same unified state of consciousness and should therefore not be considered independently.

**We did not find differences in cluster contents between the two groups. The clusters described phenomenologically distinct experience states**. To explore this idea further, we considered each dimension as a mutually dependent description of an experiential state space. To represent this in our analysis we considered the twelve dimensions to be describing a unified phenomenological state that could change dynamically over time. Using k-means clustering we identified four experientially meaningful clusters (**Figure 3**), which differ significantly in their characteristics across several experience dimensions.

**Figure 3.**
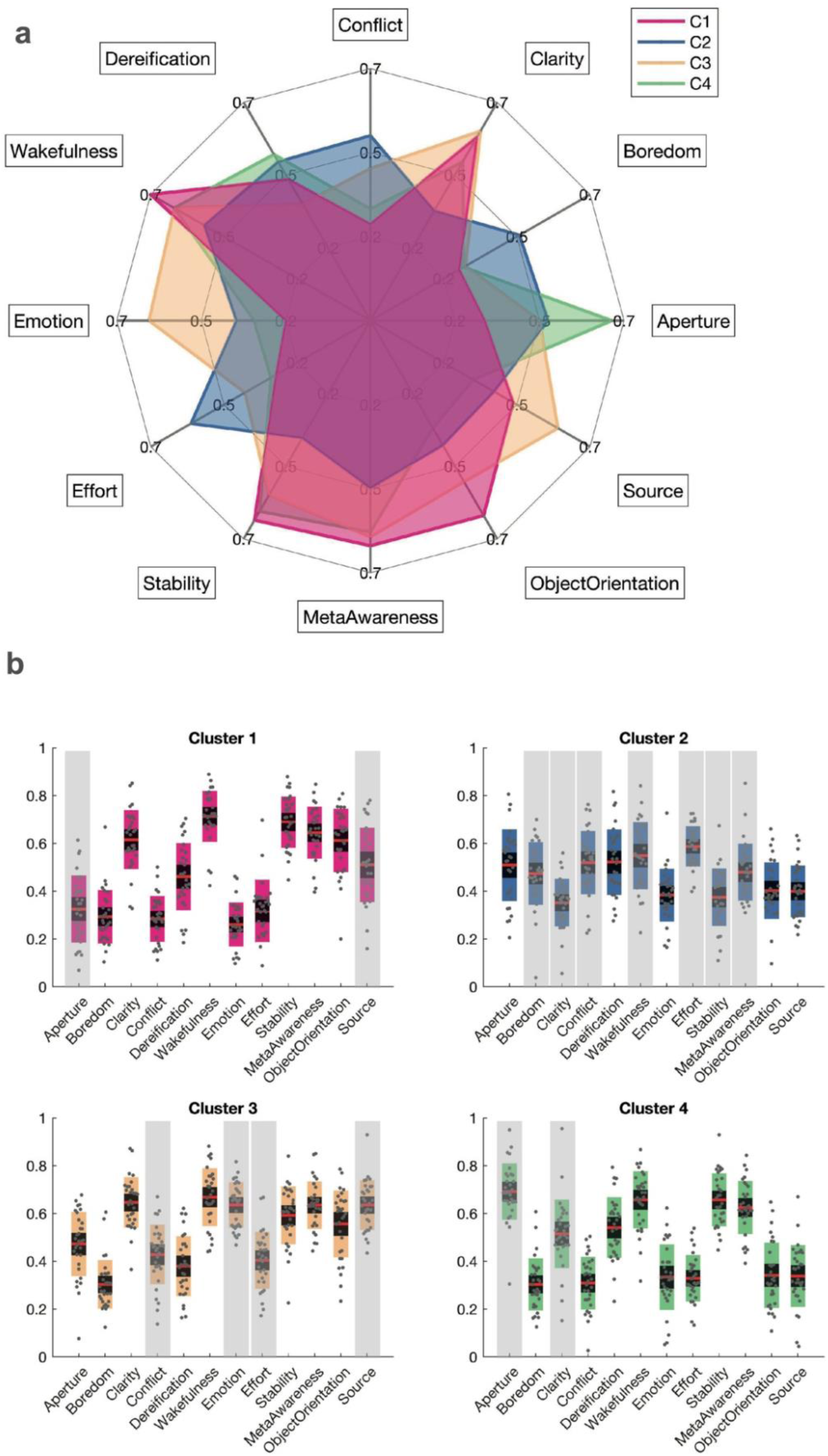
**a** Radar plot of the four cluster centroids shows that the clusters capture moments of differing mean experience dimension intensities. **b** for each cluster, the centroids were recomputed for each participant to illustrate the variance of data points included from different participants in each cluster. For each dimension a 2-way ANOVA was conducted with the factors “Group” and “cluster Identity”. The asterisk above a dimension indicates that the dimension was reliably different in this cluster from the same dimension in all the other clusters (p<.05 after Benjamini Hochburg correction). Red line is the data mean, black shading is the 95% Confidence Interval, and the coloured shading represents one standard deviation from the mean.

To characterise each cluster, we recomputed the centroid values on the participant level (**Figure 3b**). The centroids are computed as the arithmetic means of each feature (i.e., experience dimension) within a cluster. In a two-way ANOVA, the effects of Group (Home or Retreat) and cluster (clusters 1 to 4) on the intensity of each subjective experience dimension were tested. After correction we did not find a reliable interaction between the factors ‘group’ and ‘cluster’, suggesting that the effect of cluster identity on each dimension was similar in the two groups. This indicates that the clustering algorithm was probably able to group similar data points from both meditation groups into the same cluster. There was a reliable main effect of the factor ‘Group’ on two of the twelve dimensions, Wakefulness and Object Orientation, reflecting on average higher reported values of Wakefulness and Object Orientation in the Home meditators (**Figure 2**).

The factor ‘cluster’ had a reliable effect on all of the dimensions after correction for multiple comparison at a significance level p<.0001 (‘Aperture’, *F(1,3)= 46*.*1, η*^*2*^_*p*_*=*.*62*; ‘Boredom’, *F(1,3)= 21*.*96, η*^*2*^_*p*_*= 0*.*44;* ‘Clarity’, *F(1,3)= 43*.*98, η*^*2*^_*p*_ *= 0*.*61;* ‘Conflict’, *F(1,3)= 37*.*49, η*^*2*^_*p*_ *=. 0*.*57*; ‘Dereification’, *F(1,3)= 16*.*25, η*^*2*^_*p*_*= 0*.*37*; ‘Wakefulness’, *F(1,3)= 15*.*86, η*^*2*^_*p*_*=*.*36*; ‘Emotion’, *F(1,3)= 80*.*47, η*^*2*^_*p*_*=*.*74*; ‘Effort’, *F(1,3)= 69*.*73, η*^*2*^_*p*_*=*.*71*; ‘Stability’, *F(1,3)= 79*.*88, η*^*2*^_*p*_ *=*.*74*; ‘Meta-Awareness’, *F(1,3)= 30*.*50, η*^*2*^_*p*_*=*.*52*; ‘Object Orientation’, *F(1,3)= 43*.*98, η*^*2*^_*p*_*=*.*61*; ‘Source’, *F(1,3)= 37*.*14, η*^*2*^_*p*_ *=*.*57)*. This highlights that each dimension contributed to the differentiation of the centroids of the four clusters. It also indicates that each cluster had different characteristics, creating the impression of phenomenologically distinct states described by the participants through the traces.

### The cluster contents reflect three distinct ‘on-task’ and a common ‘off-task’ state

To understand in which dimensions the clusters differed, we conducted a post-hoc multiple comparison, correcting the results using Tukey’s Honestly Significant Difference Procedure (the pairwise mean differences in dimensions between clusters are presented in the **Supplementary Table 1**). The asterisks in **Figure 2** indicate the dimensions in a cluster, which were reliably different from the same dimension in the other clusters.

This allows us to examine the contents of the clusters further. Cluster 1 had reliably lower Aperture values than all other clusters, in line with a state of focused attention with a narrow spotlight of attention. It also had reliably higher values of Source compared to clusters 4 and 2, but reliably lower than cluster 3, indicating that participants perceived to be actively generating the contents of consciousness.

Experience cluster 3, which was most frequent in the Loving Kindness meditation (**Figure 4**) had reliably higher Emotion and Source values than all other clusters, and reliably higher intensities of Effort and Conflict than in cluster 1 and cluster 4, but reliably lower Effort and Conflict than captured in cluster 2. This is reminiscent of the state participants are aiming for in Loving Kindness meditation, where the instructions are aimed at the cultivation of love for all beings. Conflict can arise during this practice when the positive affective feelings are sent to a person with whom one has a difficult relationship^59,60^.

**Figure 4.**
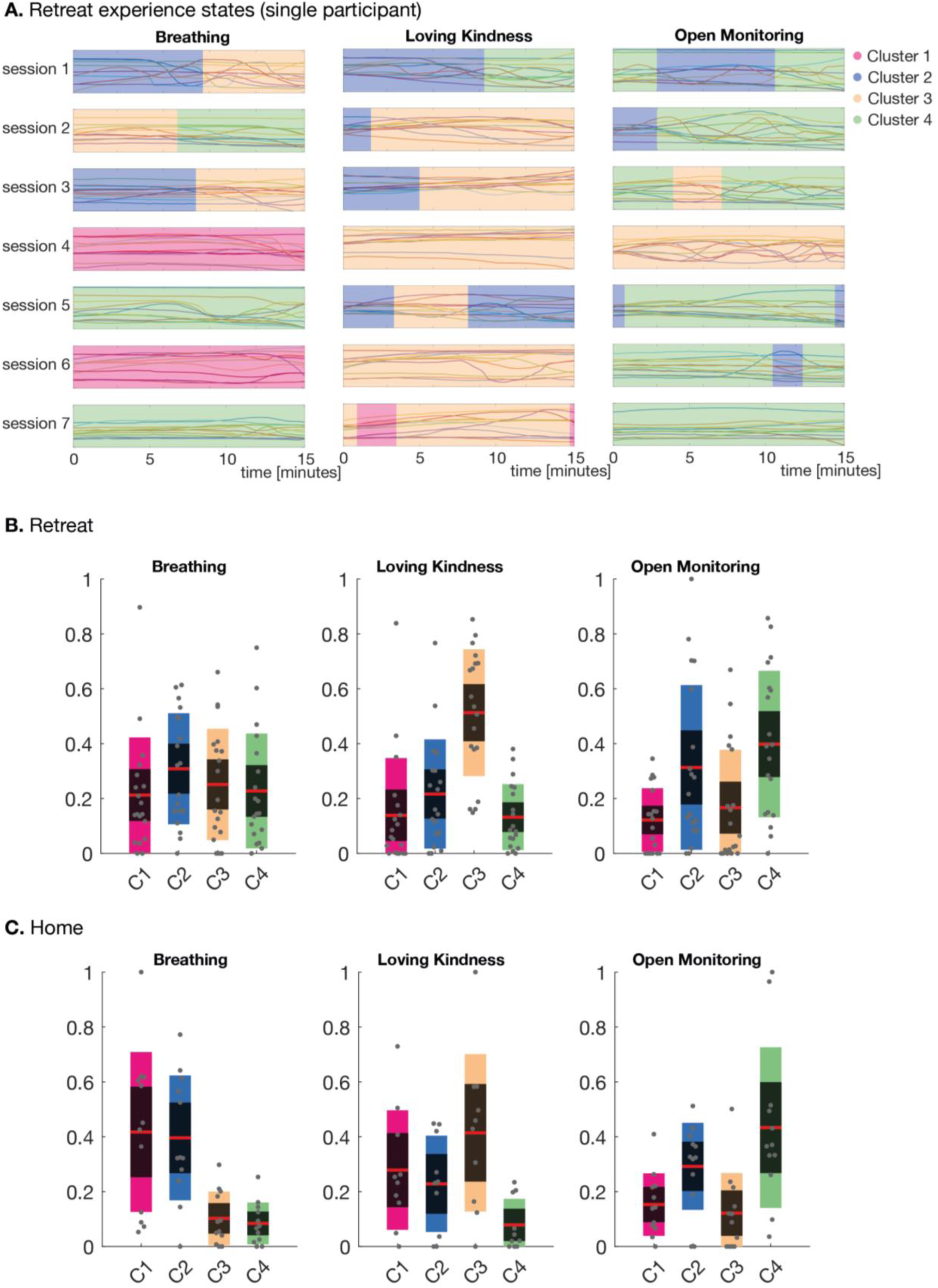
The distribution of clusters per meditation style show similarities between the two participant groups. **A**. The twelve experience traces of a single participant (Retreat) are shown for all seven sessions in each meditation style. The background shading represents the cluster attribution of each data point in the time series. **B. and C**. the relative frequency of each cluster during a meditation practice depended largely on the meditation style practiced. Cluster distribution is largely in agreement with the targeted content of the meditation style. The ‘off-task’ cluster 2, was significantly less common in Loving Kindness meditations than in Breathing meditations.

Experience cluster 4 can be differentiated by higher Aperture values than all other clusters. This cluster was most frequent during Open Monitoring meditation (**Figure 4**) and may reflect a state with a wide spotlight of attention, a key aim in OM meditation. Interestingly, cluster 4 was associated with lower levels of Clarity than cluster 1 and cluster 3, which were the states associated with Focused Attention and Loving Kindness meditation respectively. The levels of Clarity were however still higher than in cluster 2.

Cluster 2 had reliably higher values of Boredom, Conflict and Effort than all other clusters, and reliably lower values of Clarity, Wakefulness, Stability and Meta-Awareness than all other clusters. Since the dimensions this cluster scores highly on are associated with negative affect (Boredom, Conflict and Effort) and since cluster 2 does not describe any of the meditation style-specific contents, we will refer to this cluster in the remaining text as the “cluster of difficulties”. This is merely for ease of discussion and is not intended to make a claim on the causes of this cluster. Cluster 2 was less frequent overall in the Retreat Participants (*M=*.*28; SD=*.*24*) than the Home Participants (*M=*.*31; SD=*.*2)* suggesting that the participants on average experienced fewer periods of “difficulties” while meditating in the retreat setting. Cluster 2 was seen less frequently in Loving Kindness meditation (*Retreat: M=*.*22, SD= .20* ; *Home: M=*.*23, SD=*.*18)* than in Breathing meditation (*Retreat: M=*.*31, SD= .20* ; *Home: M=*.*40, SD=*.*23)* and Open Monitoring meditation *(Retreat: M=*.*31, SD=*.*30* ; *Home: M=*.*29, SD=*.*16*). Temporal experience tracing was therefore able to capture the experiences commonly associated with each meditation practice, while also capturing various off-task experiences.

### Meditation style practised impacts the time spent in each experience cluster

Supporting the observation that three experience clusters captured the target states of the three meditation styles, these experience clusters were relatively more likely to occur while the respective meditation style was practised. This result is important as the experience clusters are data-driven reconstructions from the traces of each session and not explicitly defined by the experimenters or participants.

The fraction of time spent in each cluster over the course of a meditation session depended to a large degree on the meditation style practised (**Figure 4B+C**). Repeated-measures ANOVA (within-participant factors: ‘meditation style’, ‘cluster’; between-participant factors: ‘group’) did not find a reliable main effect of group, cluster, or meditation style, nor any interaction effects related to the group factor, on the relative time spent in each cluster. There was however a reliable interaction between meditation style and cluster (*F(6) = 18*.*659, p < 0*.*001*). We did not find a three-way interaction between Group, Style and Cluster. This suggests that the distribution of clusters across a meditation session was largely determined by the meditation style practiced. Post-hoc analysis demonstrated that the clusters related to the meditation style practiced were relatively more frequent than the other clusters. This was the case in all meditation styles. Interestingly, Cluster 2 (the ‘off-task’ cluster) was significantly more common in the Breathing meditation practice than in Loving Kindness meditation (**Supplementary Table S2**). This may hint to a potential ‘ease’ of Loving Kindness meditation that may contribute to participants staying ‘on-task’ for longer periods of time.

### The relative frequency of each cluster changed as a function of time revealing style-specific dynamics shared within a group

To quantify the dynamics of the metastable states, and represent the common temporal dynamics between participants, we examined the mean relative frequency of each cluster across participants as a function of time. The area graphs in **Figure 5** show that the frequencies with which each cluster occurred in a meditation were not evenly distributed over time. Cluster 3, which was more frequent in Loving Kindness meditations (**Figure 4 A+B**), increased in frequency over the course of the meditation in both Groups. The same is true for cluster 4, which increased over the course of Open Monitoring meditations in the home meditator group. These changes of frequencies suggest that some transitions between clusters, or metastable states, were more likely than others – depending on the meditation style and group. This led us to examine transition likelihoods using Simple Markov Chain Models.

**Figure 5.**
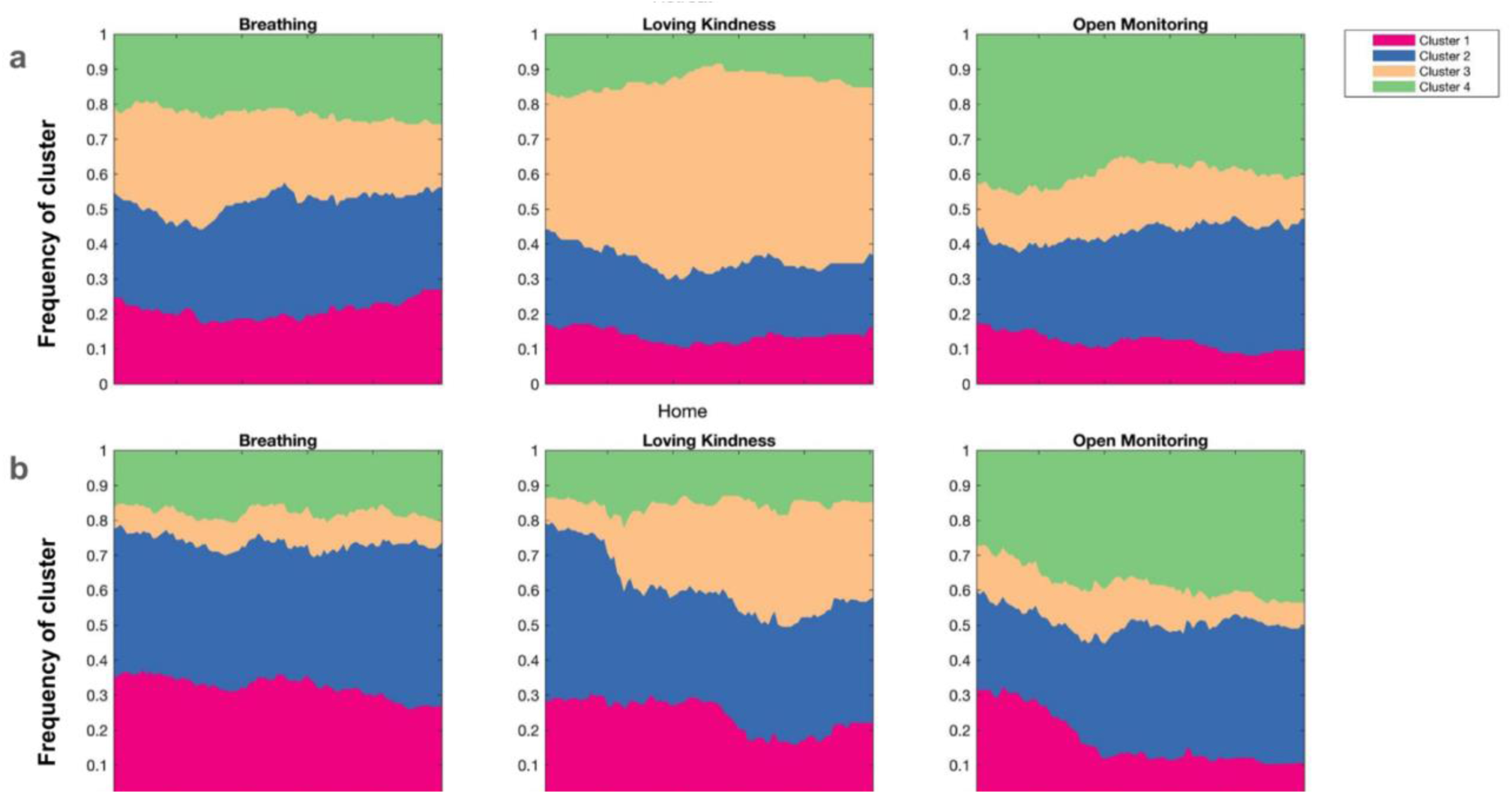
The temporal dynamics of the experience clusters per meditative state. The area plots represent the stacked relative frequency of each cluster averaged across participants for each timepoint, displayed for each meditation style and participant group. Temporal dynamics of the clusters become apparent, such as a steady increase of cluster 3 appearance during Loving Kindness meditation in both groups, a steady increase over time in cluster 4 frequency in Home Open Monitoring meditation, and an increase in cluster 2 in both groups during Open Monitoring Meditation. Cluster 2 appears to give way to cluster 3 in the Home Loving Kindness Meditation.

### Transition frequency was higher in the Home practitioners than Retreat participants

Over the 852 analysed meditations, 1385 transitions took place, with a mean of 1.47 transitions per meditation. Transitions were more frequent in the Home meditation Group (*m = 1*.*76; SD = .81 transitions per meditation*) than in the Retreat Group (*m = 1*.*19; SD = .61 transitions per meditation*). This difference was not fully accounted for by session duration, as Retreat participants transitioned 0.08 times per minute (*SD = .041*), compared to Home Group participants who transitioned 0.1 times per minute (*SD = .045*). In the Retreat group, fewer transitions per minute were made during the Loving Kindness Meditation (*m = .061 SD = .030*) than during Breathing (*m = .091 SD = .058*) and Open Monitoring (*m = .085 SD = .067*). In the Home meditation group cluster transition frequency was similar between the three meditation styles (*Breathing: m = 0*.*097 SD = 0*.*06; Loving Kindness m = 0*.*096 SD = 0*.*08; Open Monitoring: m = 0*.*106 SD = 0*.*06*). The likelihood of transitioning into the same state as the current state was very high (>.95) in all conditions, which shows the high stability of the states. The average cluster dwell time across all participants was 6.77 minutes (*SD = 5*.*83 minutes*).

For each meditation style and group, we averaged the transition matrices of all participants. From these averaged transition matrices, we computed Simple Markov Chain models to examine the transition likelihoods between the four clusters (**Figure 6**). Using these models, we subsequently ran 100-step simulations from even starting distributions (likelihood of .25 for each cluster) (e.g., **Figure 6B**), removing the influence the starting distribution had on the likelihood of each cluster during the remaining meditation.

**Figure 6.**
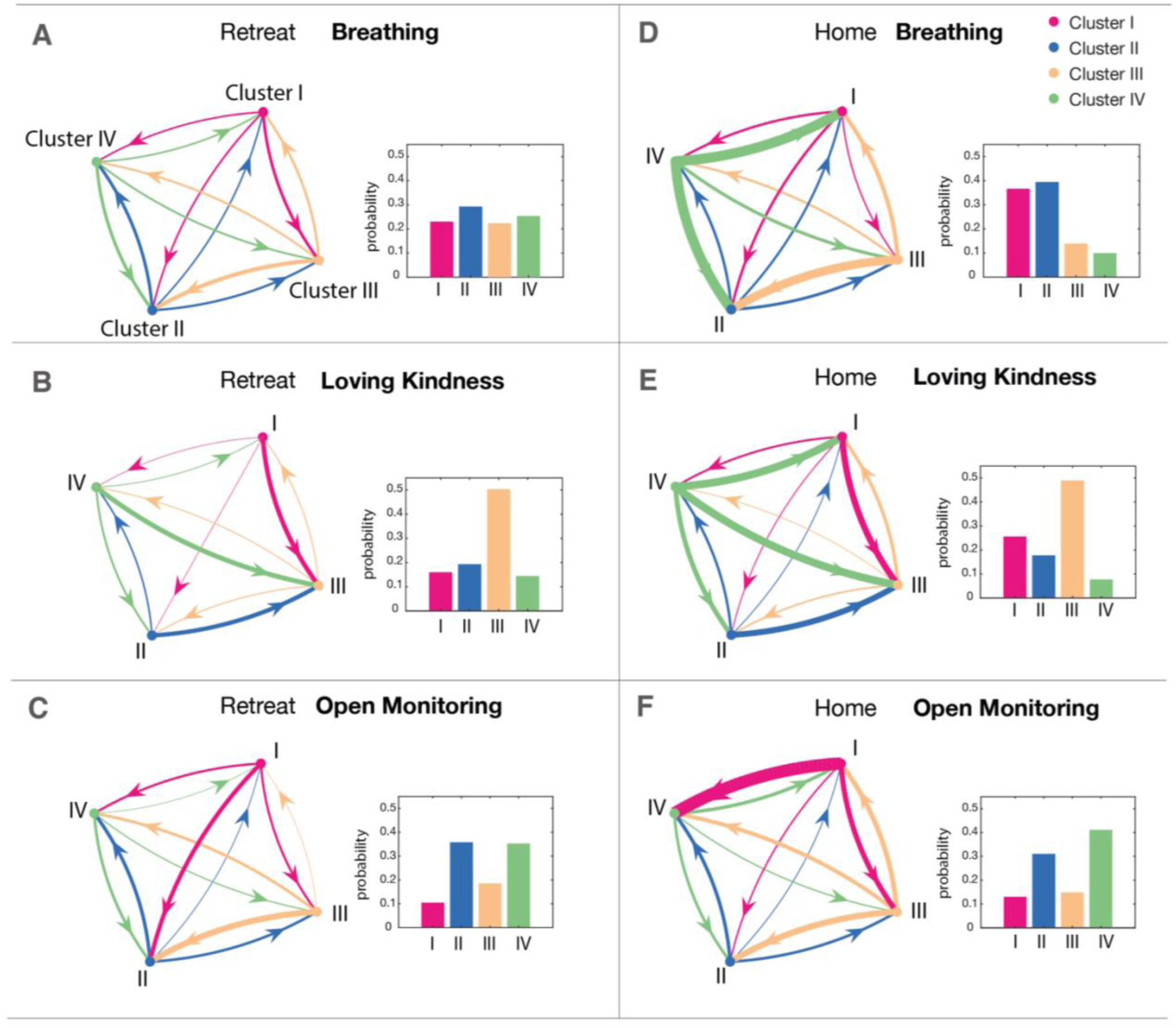
The Markov Chains show the transition likelihoods across meditation styles with some similarities between groups. The bar plots to the right of each Markov Chain show the evolution from a uniform distribution after 100 steps (equivalent to the duration of one meditation session) in the Markov Chain model. The width of the edges represents the relative likelihood of the transition. Markov Chain graphs exclude self-transitions. **A**,**B**,**C** show the Markov Chains and simulated distributions for the retreat group, **D**,**E**,**F** the Markov Chains and simulated distributions for the home group. In Loving Kindness meditation in both groups, the simulation shows an increased likelihood of cluster 3. In Open Monitoring meditation, the simulations show an increased likelihood of cluster 2 and cluster 4. This corroborates the empirical findings presented in Figure 4. This simulation demonstrates that the uneven starting distribution in the empirical data is not required for generating the observed trends in state distributions between the meditation styles.

The simulations revealed some interesting properties of the systems. The simulated end distributions were strikingly similar to the empirical mean distributions (**Figure 4B+C**), despite the difference in simulated vs empirical starting distributions. Further, the model and simulations showed a clear elevation in the probability of one or two clusters in each meditation style, with agreement in both Groups. An exception to this was the Retreat Group practising Breathing meditation, where there was a striking lack of a dominant cluster after 100 simulation steps.

Three of the four clusters are highly prevalent in one meditation style only. Cluster 4 was highly likely to be transitioned into during Open Monitoring meditation. Cluster 3 was highly likely to be transitioned into during Loving Kindness meditation, and less likely transitioned out of, equally creating an increased accumulation in cluster 3. The transition into cluster 1 was highly likely in Home-meditators during Breathing meditation.

Cluster 2 was the only cluster that has a high likelihood after 100 steps in more than one meditation style. The frequency of transition into cluster 2 was high in Breathing and Open Monitoring meditation, in both groups. Noteworthy might be that the likelihood of a participant being in cluster 2 after 100 simulation steps was markedly lower in Loving Kindness meditation than the other meditation styles and that this was true for both groups.

## Discussion

In this study we used temporal experience tracing to capture the dynamics of experiences in two groups of meditators practicing three different meditation styles in different contexts. The advances made here are as much to do with meditation as they are to capturing the experience dynamics of any task that humans may do, practice or endure. We demonstrate that the temporal dimensions– allowing participants to express how the dimensions changed in intensity over time – captures the dynamic interactions between the dimensions, in the form of meaningful, recurring patterns of common experiences. We captured these patterns in the form of clusters using a fully data-driven approach, *k-means* clustering. These data-driven clusters were metastable states of reconstructed experience, where participants transitioned on average 1.47 times per meditation (every 6-7 minutes). Three of the four data-driven clusters reflected metastable experience states relating to the intention of the meditation practice, and the remaining cluster of Experience (cluster 2) captured a low motivational state, characterised by high Boredom, Conflict and Effort, low Wakefulness, Clarity and Stability, and was equally common in both meditation groups. Finally, we were able to characterise the transition likelihood between clusters through Simple Markov Chains, simulating the frequency of each cluster after 100 timepoints in a scenario where the starting distributions were even across clusters. This simulation showed that participants were, for example, less likely to transition into cluster 2 during Loving Kindness meditation than while practising other meditation styles. It also revealed commonalities in the transitions between groups and meditation styles in convergence to the other parameters assessed in the search of the dynamics of experience. These results show that the collection of continuous experience data can lead to a much better understanding of the dynamics of experiences as perceived by the participant, and the results represent an advance in the treatment of phenomenology as dynamical intensities of experience.

How we experience events is a key aspect of science as it represents directly how we think, and how we feel, and ultimately, who we are^61^. One way to describe our experiences is through time, how they happened, what we thought, how we felt and what we did^62^. This concept of dynamics of experience is what we aim to formalize in this work, the quantification of the Stream of consciousness, a concept developed in many forms throughout history^63^. In particular, and relevant for behavioural sciences, is the work by William James, who defined that even the “most dim shade of perception enters into, and in some infinitesimal degree modifies, the whole existing state”^17^, setting the scene to, a hundred years later, integrate cognitive neuroscience with the study of human experience^64^. We subscribe to this view of integrated conscious experience, and managed to reconstruct in this work a data-driven state built from the time-continuous traces of meaningful aspects of experience. This is by no means a complete description of the mental state and flow of thought of our participants, but a reductive representation of the stream of thought that allows for a flexible and creative analytical approach for cognitive neuroscience. The view of an integrated conscious experience in the stream is not without its detractors as Strawson suggests that “our thought is fluctuating, uncertain, fleeting […] It keeps slipping from the mere consciousness into self-consciousness and out again”, and this is especially so when one is just sitting and thinking^65^. If another framework, one with non-contiguous, interrupted stream is taken, it will be possible to make other assumptions to the data collected in this project and reshape the hypotheses and analyses to further test theories of dynamics of consciousness that support interrupted streams of consciousness.

The approach we took to deal with the variance in experiences and content of thought, was to define – in agreement with the experts in those experiences – the meaningful phenomenological features we called dimensions, in meditation. This allowed for a reduction in the complexity of reporting and for common ground between participants and sessions. By developing the Temporal Experience Traces method, we added the passage of time to capture the stream of consciousness, increasing the data dimensionality. In the need to reduce the complexity and approximate the metastable states of experience in the flow of thought, we reconstructed the states via k-means clusters. Others have used, for example, principal components analysis to reveal the latent structure of thought on experience sampling experiments of mind-wandering^66,67^, but this differs from our results as their method is not applied on a stream of dynamical manner but to 9 questions using a 9-point Likert scale sampled intermittently during a task^68^. This approach of data reduction and finding common aspects of the content of consciousness has been applied in various forms before and the key differences from the current method in the conceptual advance of using time series to represent important aspects of experiences and reconstruct metastable states from this multifactorial representation of the stream of consciousness.

The k-means algorithm allowed to partition the 852 sessions, 12 traces of dimensions each, into distinct non-overlapping subgroups, the metastable clusters of experience. In the past, the k-means algorithm has been frequently used to identify clusters, or states in resting data fMRI, as well as EEG data^6–8^. By applying the algorithm to continuous subjective experience data, we capture reconstructed experience states from the perspective of the experiencer (first person). An alternative approach to this would be the use of a Hidden Markov Model (HMM), which would determine both the hidden structure (or clusters) of the data and their transition likelihoods. For example, Cabral et al compared results from k-means clustering to the HMM in research on cognitive performance of patients and found them to be similar^69^. Here we opted for the k-means algorithm due to its simple implementation and low computational cost. More importantly, both theoretically and methodologically, capturing the dimensions of experience in time series (through TET) and using pattern extraction and data reduction methods allows for the phenomenology to approximate cognitive imaging data, putting them on common ground for signal-level processing, with the need to further bridge them conceptually but maybe not methodologically.

In this exploration of the data-driven reconstructed states we interpreted three of the four clusters qualitatively matching specific mental states aimed in each of the meditation styles. In the theoretical framework proposed by Lutz, from where we derived some of the dimensions used in this study, Breathing meditation is described as a style of focused attention meditation, where experienced meditators could approach a state of high Clarity, low Aperture, high Stability, low Effort, high Dereification and high Object Orientation. These features were present in our cluster of Experience 1, which was most frequent in Home meditators during Breathing and Loving Kindness meditation. It is noteworthy that cluster 1 was not common in Retreat participants, who had less experience with meditation. Cluster 1 was consistently the least likely to be transitioned into from cluster 2, the cluster of difficulties. The Euclidean distance between the centroids of clusters 2 and 1 was also the highest. We therefore hypothesise that cluster 1, a state of focused attention, is the most difficult to attain for participants who are lingering in a metastable state of difficulties, or low motivation (cluster of Experience 2).

This cluster of difficulties, cluster of Experience 2, was dynamically marked by high levels of Boredom, Conflict and Effort, and low levels of Clarity, Wakefulness, Stability and Meta Awareness. This cluster was present in all meditation styles, in both Groups, unsurprisingly. The experiences this cluster appears to represent, are often described in meditation teachings as “hindrances” (for example Joseph Goldstein & Jack Kornfield 2001 in “Seeking the Heart of Wisdom” name irritation, boredom, fear, doubt and restlessness as common hindrances), and confirmed with participants in subsequent conversations after the identification of the Clusters of Experiences. In the simulations derived from the Markov Chains, Breathing and Open Monitoring meditation led to high likelihoods of cluster 2 occurring. Interestingly, both meditation groups found themselves less frequently in cluster 2 during Loving Kindness meditation than any of the other meditation styles. Participants were also less likely to transition into cluster 2 during Loving Kindness meditation, and more likely to transition away from cluster 2 during Loving Kindness meditation. This finding is in line with the understanding that boredom arises from a state of high effort in “uninteresting tasks” and is encouraged by lower levels of alertness^44,45^. The subjective experience of cognitive effort is thought to be a potential mechanism for tracking the cost of working memory allocation to a particular object of attention, and may be both tracked and enabled by phasic and tonic Dopamine levels^48^. It is therefore likely that Loving Kindness meditation, a style of meditation that encourages active generation of emotions and visual imagery, offers more complex content for cognitive processing and is inherently more engaging than focused attention meditation and Open Monitoring meditation. If more engagement lead to ease to perform, motivation would increase and the likelihood of cluster of difficulties (cluster 2) arising would be lower.

The reverse inference of the composition of the clusters of Experience allow for a meaningful – perhaps parsimonious – interpretation of all reconstructed states, at least to the degree that can be trusted as a basis for theory building and exploration^70,71^. For cluster of Experience 3 it became apparent that it was most common in both groups during Loving Kindness meditation and that it was characterised by low Conflict and Dereification, and high Emotion and Source, thereby matching the expected aspects of a successful experience of Loving Kindness meditation. The high values in Source, which reflects how active, or generative the practice is, are coherent with the visualisations and active generation of emotions commonly practiced in Loving Kindness meditation. Retreat participants were more likely than Home meditation practitioners to start the Loving Kindness meditation in cluster 3, which may be a reflection of the contextualisation and intention setting by the meditation teacher on the retreat and led to practitioners finding themselves in the cluster as soon as they started the meditation. Even though Retreat participants were likely to start their meditation in cluster 3, it was also the cluster most frequently transitioned into from other clusters during Loving Kindness meditation, in both Groups. This was highlighted by our simulations, in which we used the Markov chain models on an even starting cluster distribution, resulting in a high likelihood of cluster 3 after 100 steps (corresponding to the length of one meditation), especially for the Home group.

In the original framework, Lutz and collaborators^27^ set out the phenomenological matrix of meditative states and hypothesised that Open Monitoring meditations, when conducted by experienced meditators, would be accompanied by high Aperture, Clarity, Stability, Dereification and Meta-Awareness, as well as low Effort and Object Orientation. This description fits well with the content of cluster 4, which had reliably higher Aperture ratings and reliably lower Object Orientation than the other clusters. The levels of Clarity, Stability, Dereification, Meta-Awareness and Effort were similar to the levels achieved in the clusters 3 (Affective cluster) and 1 (Focused Attention cluster). In both meditation groups cluster 4 was most common in Open Monitoring meditation, and Home group participants were likely to transition into cluster 4 in the course of the open monitoring meditation. Cluster of Experience 4, as an algorithmically reconstructed state that we interpret as a metastable state associated with the intended experience of Open monitoring, it differentiates from the other 3 states mathematically and descriptively but also partially converges with the phenomenological description by Lutz^27,72^ despite the different methodology applied to characterise the contents of the mind in meditation.

The granularity of experiences picked up by the clusters, inadvertently depends on the number of clusters chosen *a priori* for the clustering algorithm. The higher the number of clusters (k) chosen, the better the fit of the cluster centroids describing the data. The best fit in k-means analysis therefore will always be found at a k equal to the number of datapoints itself, but the result would be meaningless. We chose 4 clusters based on the ‘elbow method’, as the point where the difference in error obtained with the next higher k plateaued. Using four clusters happened to capture the main aspects of each meditation style, as well as a cluster of disengagement from the tasks; however, it is not inconceivable that an increased number of clusters would lead to a more detailed picture and might capture some differences between the two meditation groups, or differentiate between different aspects of ‘difficulties’, which are currently all present in a single cluster. It is interesting to note however, that the differences between meditation styles were greater than differences between two completely separate meditation groups with different levels of experience, leading to the clustering algorithm to partition the data by experiences associated with the meditation styles, rather than the level of expertise of participants. This result is encouraging as it highlights the convergence of the traces of experience as a valid tool in 2 independent groups of people and should inspire further research in task-related psychology in conjunction with behavioural, computational and imaging methods.

This study presents a powerful way to capture the temporal dynamics of subjective experiences of continuous states. While mindfulness meditation research was a natural starting point for the application of this technique, with an established basis of its phenomenological structure, this technique may be of relevance to other areas of research, such as pain research, agency research, in altered states of consciousness research, or in other categories of meditation not primarily involving attention or monitoring^73^. If applied to these cases, researchers might consider an empirical approach to the phenomenological ‘front-loading’, by conducting phenomenological interviews on a subsample of participants. This might help develop the phenomenological structure, or state-space relevant to the study.

While we have not presented any neurophysiological data, and the temporal granularity of the states is limited by the physical parameters of the hand-drawn graph, the question arises how the clusters of Experience may relate to physiological mechanisms and timescales. For example, some research suggests that discrete neuromodulator pathways drive Focused Attention and Open Monitoring states^74^. Focused Attention may be primarily regulated by dopaminergic pathways, whereas Open Monitoring practices were suggested to be primarily driven by noradrenergic pathways^74,75^. Combining this research with a method that captures the perceived temporal dynamics of the experiences associated with each practice, future research may generate a better understanding for the (top-down) transitions into the neurophysiological states that may be required in order to generate and stabilise the desired phenomenological experience states.

To conclude, using temporal experience tracing can paint a detailed picture of the phenomenological dynamics of continuous tasks. It tells us that both of our Groups, even though they had different levels of experience with meditation, went through periods of subjectively difficulty, or ‘off-task’ time (cluster 2), while alternating between metastable states of experience. This may be an important consideration when examining the neurophysiological correlates of meditation states, as these periods of difficulty may dilute the differences between meditation states, if they are parsed by the meditation style only. We hope that from a practical point of view the traces of experience can help modulate, conceptualise and improve the measurements and understanding of cognition by taking into account the stream of thought of the participants.

## Acknowledgments

ESRC Fellowships and Mind and Life Foundation European Varela Award to BJ; Fieldwork Grant, Department of psychology, University of Cambridge.

