## Supplementary Information for "Drawing the experience dynamics of meditation"

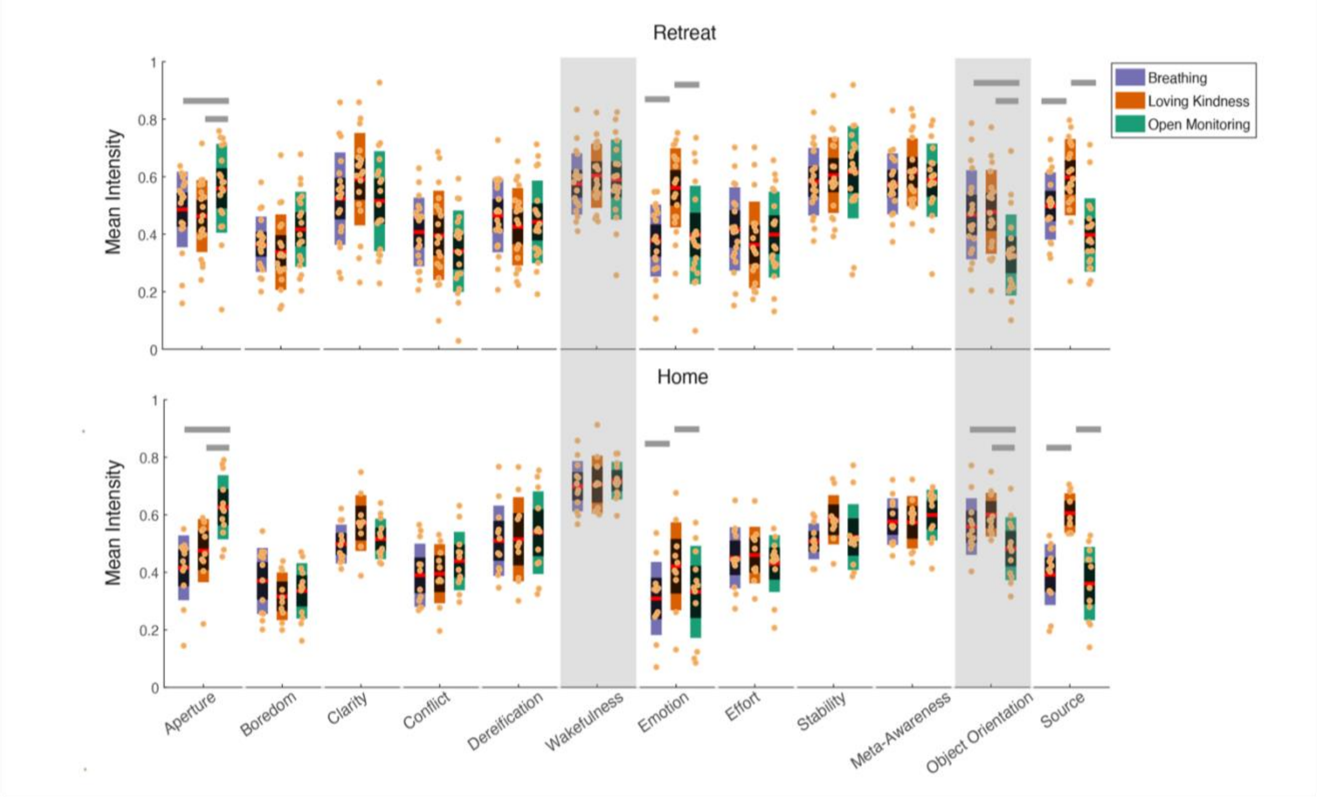


***Figure S1****: Mean dimension intensity values for each participant separated by meditation style show widespread agreement between the two groups. Top: Retreat participants Bottom: Home meditation participants. Purple: Breathing Meditation, Tawny: Loving Kindness Meditation, Green: Open Monitoring Meditation. Red line indicates the mean; black shading is the 1.96 SEM (95% CI), coloured shading represents one Standard Deviation.* ***Horizontal grey bars*** *indicate a style effect (Breathing | Loving Kindness | Open Monitoring) on dimension intensity,* ***Grey boxes*** *mark a group effect (Retreat | Home) on dimension intensity. Effects shown after correction for multiple comparisons. There were no significant interactions between Group and meditation Style for any of the dimensions.*

**Averaging dimension intensities produces some differences between meditation styles and participant groups.** Typically, questionnaires are used to quantify the intensities of experiences during meditation [76–78]. Conceptually, the analysis of experience questionnaires is based on the assumption that retrospective ratings approximate the arithmetic mean of the continuous experience over time, or alternatively, that the changes in experience states over time are negligible.

We investigated if such an approach would allow us to detect differences in subjective experiences between the three different meditation styles and between the two meditation groups who practiced in different settings. To this end the mean dimension intensities were computed for each meditation session to create a single summary rating (**Figure S1**)

We examined the effect of the factors ‘Group’ (Home/Retreat) and ‘Style’ (Breathing, Loving Kindness, Open Monitoring) on the mean intensity of each dimension in a two-way ANOVA.

Participants in the Home Group reported reliably higher Wakefulness values (M = .71, SD = 0.08) than the Retreat Group (M = .59, SD =.12, F(1, 2) = 11.43, *p* (adjusted) = .014, $\eta^2_p$=.297). Object-Orientation was also higher in the Home Group (M = .54, SD = .11) than the Retreat Group (M = .42, SD = 0.16, F(1, 2) = 11.30, *p* (adjusted) = .014, $\eta^2_p$=.295). There was no reliable interaction between the factors Group and Style for any of the dimensions, indicating that the effect of Group did not mediate the Style effect. In both groups the factor Style had a statistically reliable effect on the dimensions Aperture (Breath: M = .46, SD = .13, LK: M = .47, SD = 0.12, OM: M = .59, SD = .14, F(1, 2) = 14.41, *p* (adjusted) <.0001, $\eta^2_p$=.35), Emotion (Breath: M = .35, SD = 0.13, LK: M = .51, SD = .16, OM: M = .37, SD = .17, F(1, 2) = 18.51, *p*(adjusted)<.0001, $\eta^2_p$=.41), Object-Orientation (Breath: M = .50, SD = .14, LK: M = .52, SD = 0.14, OM: M = .39, SD = .15, F(1, 2) = 11.38, *p*(adjusted)=.0002, $\eta^2_p$=.297) and Source (Breath: M = .46, SD = 0.12, LK: M = .60, SD = .11, OM: M = .38, SD = .13, F(1, 2) = 30.838, *p*(adjusted)<.0001, $\eta^2_p$ = .533). In other words, Open Monitoring meditation was associated with higher Aperture, lower Object Orientation and lower Source than the other two meditation styles, and Loving Kindness meditation was associated with higher values in Emotion and Source than the other meditation styles. Notably, this analysis was not able to find any differences in eight of twelve experience dimensions.

***Table S1*** *The results of the pairwise comparisons between clusters for each experience dimension. CI= 95% confidence interval; p= corrected using Tukey's Honestly Significant Difference Procedure.* ***Bold*** *are the Mean differences below the significance threshold of p <.01*

| | | Cluster 2 | | | Cluster 3 | | | Cluster 4 | | |
|---|---|---|---|---|---|---|---|---|---|---|
| | | Mean Difference | CI | *p* | Mean Difference | CI | *p* | Mean Difference | CI | *p* |
| **Aperture** | Cluster 1 | **-0.17** | [-0.27 -0.08] | 0.000 | **-0.14** | [-0.23 -0.05] | 0.001 | **-0.36** | [-0.45 -0.26] | 0.000 |
| | Cluster 2 | | | | 0.03 | [-0.06 0.13] | 0.777 | **-0.18** | [-0.27 -0.09] | 0.000 |
| | Cluster 3 | | | | | | | **-0.22** | [-0.31 -0.12] | 0.000 |
| **Boredom** | Cluster 1 | **-0.18** | [-0.26 -0.10] | 0.000 | -0.01 | [-0.09 0.07] | 0.987 | -0.01 | [-0.08 0.07] | 0.997 |
| | Cluster 2 | | | | **0.17** | [0.09 0.25] | 0.000 | **0.17** | [0.09 0.25] | 0.000 |
| | Cluster 3 | | | | | | | 0.00 | [-0.07 0.08] | 0.999 |
| **Clarity** | Cluster 1 | **0.25** | [0.17 0.33] | 0.000 | -0.03 | [-0.10 0.05] | 0.839 | 0.09 | [0.01 0.17] | 0.022 |
| | Cluster 2 | | | | **-0.28** | [-0.36 -0.20] | 0.000 | **-0.16** | [-0.24 -0.08] | 0.000 |
| | Cluster 3 | | | | | | | **0.11** | [0.03 0.19] | 0.002 |
| **Conflict** | Cluster 1 | **-0.24** | [-0.32 -0.16] | 0.000 | **-0.15** | [-0.23 -0.07] | 0.000 | -0.03 | [-0.11 0.05] | 0.784 |
| | Cluster 2 | | | | **0.09** | [0.01 0.17] | 0.014 | **0.21** | [0.13 0.29] | 0.000 |
| | Cluster 3 | | | | | | | **0.12** | [0.04 0.20] | 0.001 |
| **Dereification** | Cluster 1 | -0.05 | [-0.14 0.04] | 0.434 | 0.08 | [0.00 0.17] | 0.070 | -0.08 | [-0.17 0.01] | 0.074 |
| | Cluster 2 | | | | **0.13** | [0.05 0.22] | 0.001 | -0.03 | [-0.12 0.06] | 0.782 |
| | Cluster 3 | | | | | | | **-0.17** | [-0.25 -0.08] | 0.000 |
| **Wakefulness** | Cluster 1 | **0.16** | [0.09 0.24] | 0.000 | 0.04 | [-0.03 0.12] | 0.452 | 0.05 | [-0.02 0.13] | 0.283 |
| | Cluster 2 | | | | **-0.12** | [-0.20 -0.04] | 0.001 | **-0.11** | [-0.19 -0.03] | 0.002 |
| | Cluster 3 | | | | | | | 0.01 | [-0.07 0.09] | 0.989 |
| **Emotion** | Cluster 1 | **-0.12** | [-0.20 -0.05] | 0.000 | **-0.37** | [-0.44 -0.29] | 0.000 | -0.08 | [-0.15 0.00] | 0.045 |
| | Cluster 2 | | | | **-0.25** | [-0.32 -0.17] | 0.000 | 0.05 | [-0.03 0.12] | 0.326 |
| | Cluster 3 | | | | | | | **0.29** | [0.22 0.37] | 0.000 |
| **Effort** | Cluster 1 | **-0.27** | [-0.34 -0.20] | 0.000 | **-0.10** | [-0.17 -0.02] | 0.005 | -0.02 | [-0.09 0.06] | 0.941 |
| | Cluster 2 | | | | **0.17** | [0.10 0.25] | 0.000 | **0.25** | [0.18 0.33] | 0.000 |
| | Cluster 3 | | | | | | | 0.08 | [0.01 0.15] | 0.027 |
| **Stability** | Cluster 1 | **0.31** | [0.24 0.39] | 0.000 | **0.10** | [0.02 0.17] | 0.004 | 0.04 | [-0.04 0.11] | 0.581 |
| | Cluster 2 | | | | **-0.21** | [-0.29 -0.14] | 0.000 | **-0.28** | [-0.35 -0.20] | 0.000 |
| | Cluster 3 | | | | | | | -0.06 | [-0.14 0.01] | 0.126 |
| **Meta Awareness** | Cluster 1 | **0.17** | [0.10 0.25] | 0.000 | 0.02 | [-0.06 0.09] | 0.950 | 0.02 | [-0.06 0.10] | 0.892 |
| | Cluster 2 | | | | **-0.16** | [-0.23 -0.08] | 0.000 | **-0.15** | [-0.23 -0.07] | 0.000 |
| | Cluster 3 | | | | | | | 0.01 | [-0.07 0.08] | 0.998 |
| **Object Orientation** | Cluster 1 | **0.22** | [0.14 0.30] | 0.000 | 0.06 | [-0.02 0.14] | 0.239 | **0.26** | [0.18 0.35] | 0.000 |
| | Cluster 2 | | | | **-0.16** | [-0.24 -0.08] | 0.000 | 0.04 | [-0.04 0.12] | 0.493 |
| | Cluster 3 | | | | | | | **0.21** | [0.12 0.29] | 0.000 |
| **Source** | Cluster 1 | 0.10 | [0.02 0.18] | 0.014 | **-0.12** | [-0.21 -0.04] | 0.001 | **0.17** | [0.09 0.25] | 0.000 |
| | Cluster 2 | | | | **-0.22** | [-0.31 -0.14] | 0.000 | 0.07 | [-0.01 0.15] | 0.123 |
| | Cluster 3 | | | | | | | **0.29** | [0.21 0.38] | 0.000 |

***Table S2*** *The pairwise comparison of relative time spent in each cluster by meditation style. CI= 95% confidence interval; p= corrected using the Benjamini & Hochberg (1995) procedure for controlling the false discovery rate (FDR) of a family of hypothesis tests.* ***Bold*** *are the Mean differences below the significance threshold of p <.05*

| | | Loving Kindness | | | Open Monitoring | | |
|---|---|---|---|---|---|---|---|
| | | Mean Difference | CI | *p* | Mean Difference | CI | *p* |
| **Cluster 1** | Breathing | **0.16** | [0.04 0.28] | 0.017 | **0.21** | [0.12 0.30] | 0.000 |
| | Loving Kindness | | | | 0.05 | [-0.09 0.18] | 0.659 |
| **Cluster 2** | Breathing | **0.12** | [0.03 0.22] | 0.017 | 0.05 | [-0.08 0.18] | 0.659 |
| | Loving Kindness | | | | -0.07 | [-0.21 0.07] | 0.501 |
| **Cluster 3** | Breathing | **-0.31** | [-0.46 -0.15] | 0.000 | -0.07 | [-0.05 0.20] | 0.459 |
| | Loving Kindness | | | | **0.38** | [0.25 0.51] | 0.000 |
| **Cluster 4** | Breathing | 0.08 | [-002 0.18] | 0.182 | **-0.28** | [-0.42 -0.14] | 0.000 |
| | Loving Kindness | | | | **-0.36** | [-0.51 -0.21] | 0.000 |

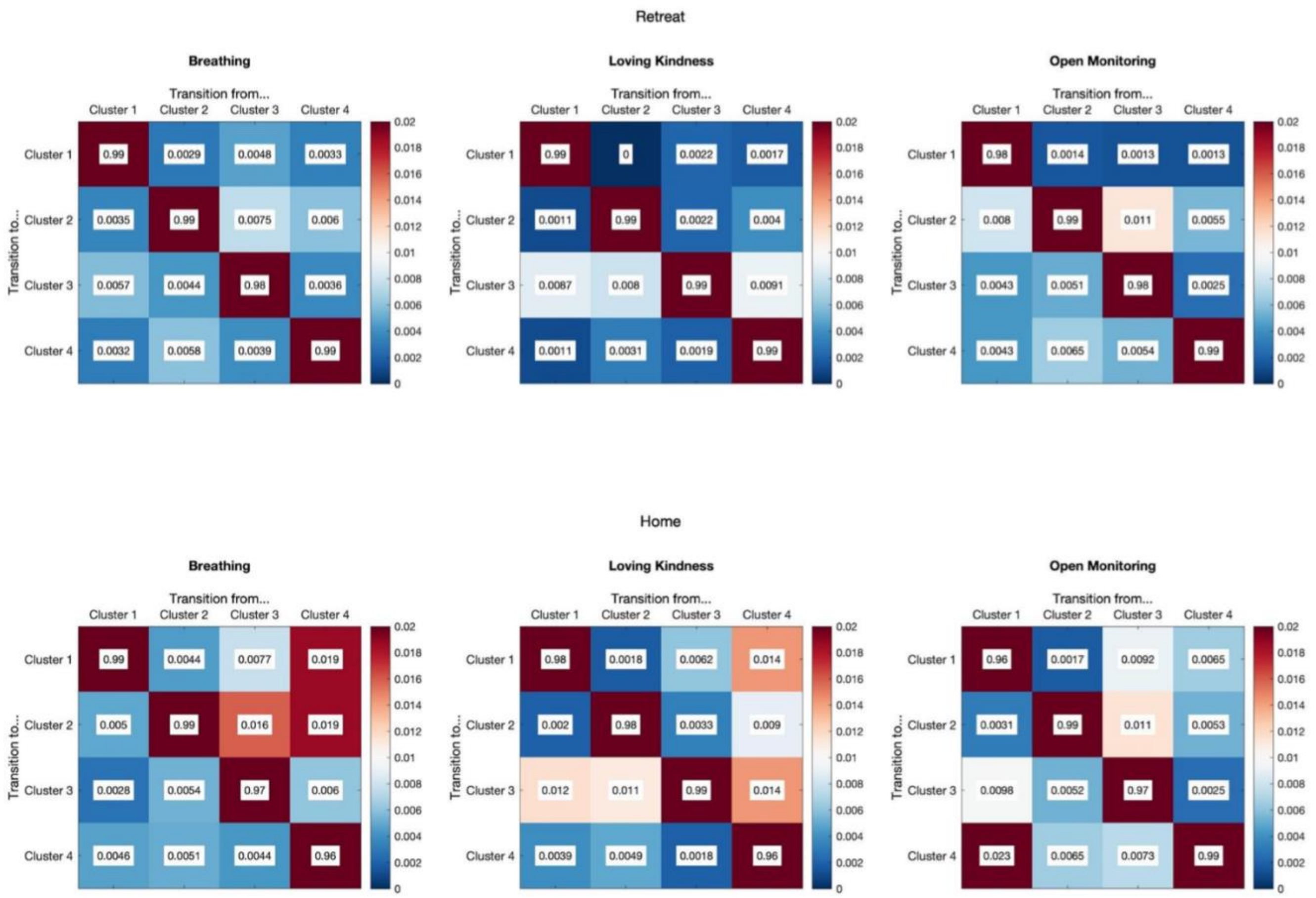


***Figure S1:*** *The transition matrices for the Retreat (top) and Home (bottom) groups show meditation-style dependent transition matrices.*

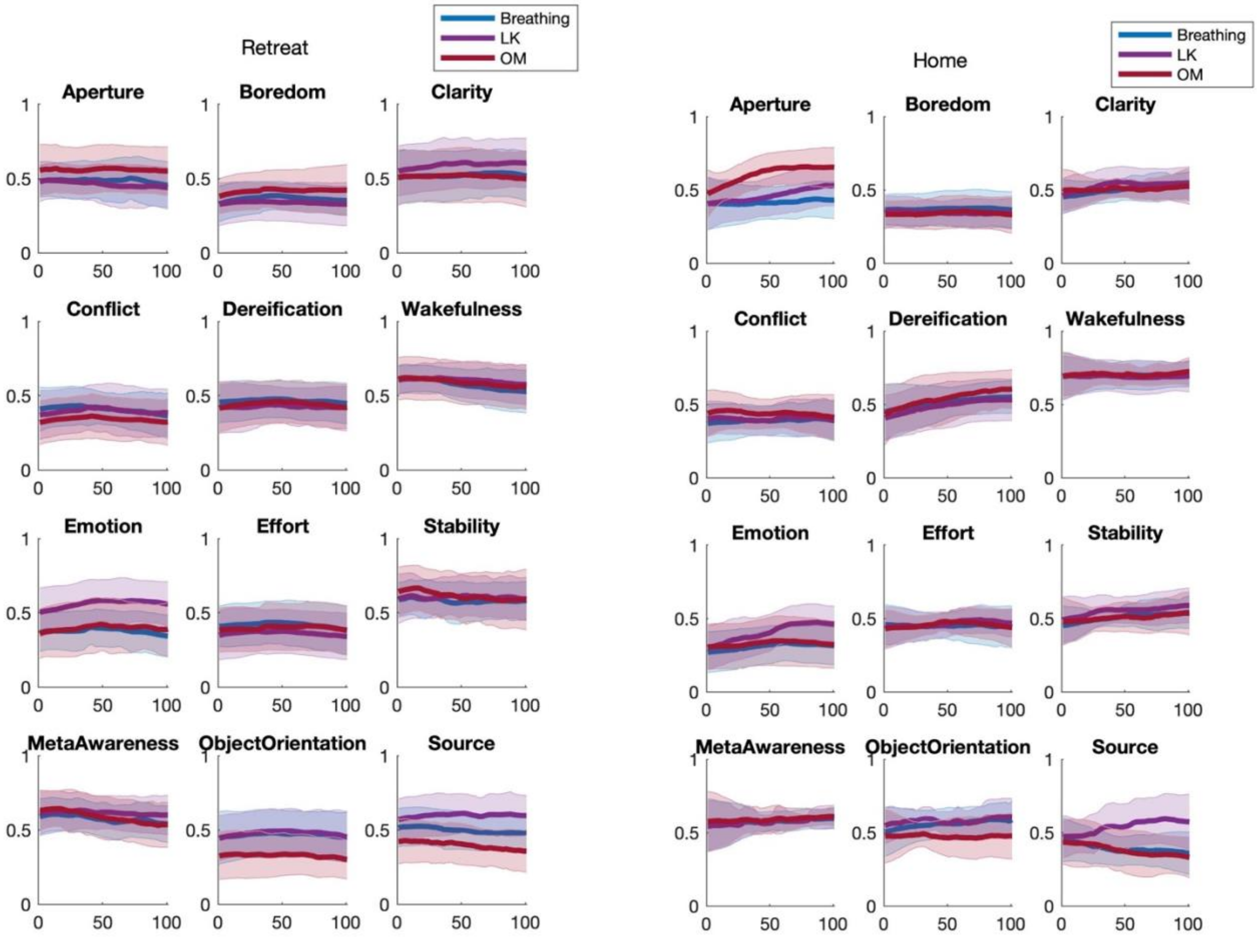


***Figure S2:*** *The group averages for each dimension over time, plotted for each meditation style, show some of the time dynamics were shared between meditation styles (e.g., Dereification and Stability in the Home meditation group), while others were not (e.g., Source, Aperture, Emotion and Object Orientation in the Home Meditation Group). There were also apparent group differences in the dynamics of some dimensions – Wakefulness dropped over time in the Retreat group across all meditation styles, while it stayed stable in the Home meditation group. This was also seen in Dereification, which increased in the Home meditation group for all meditation styles, while it stayed stable in the Retreat group.*

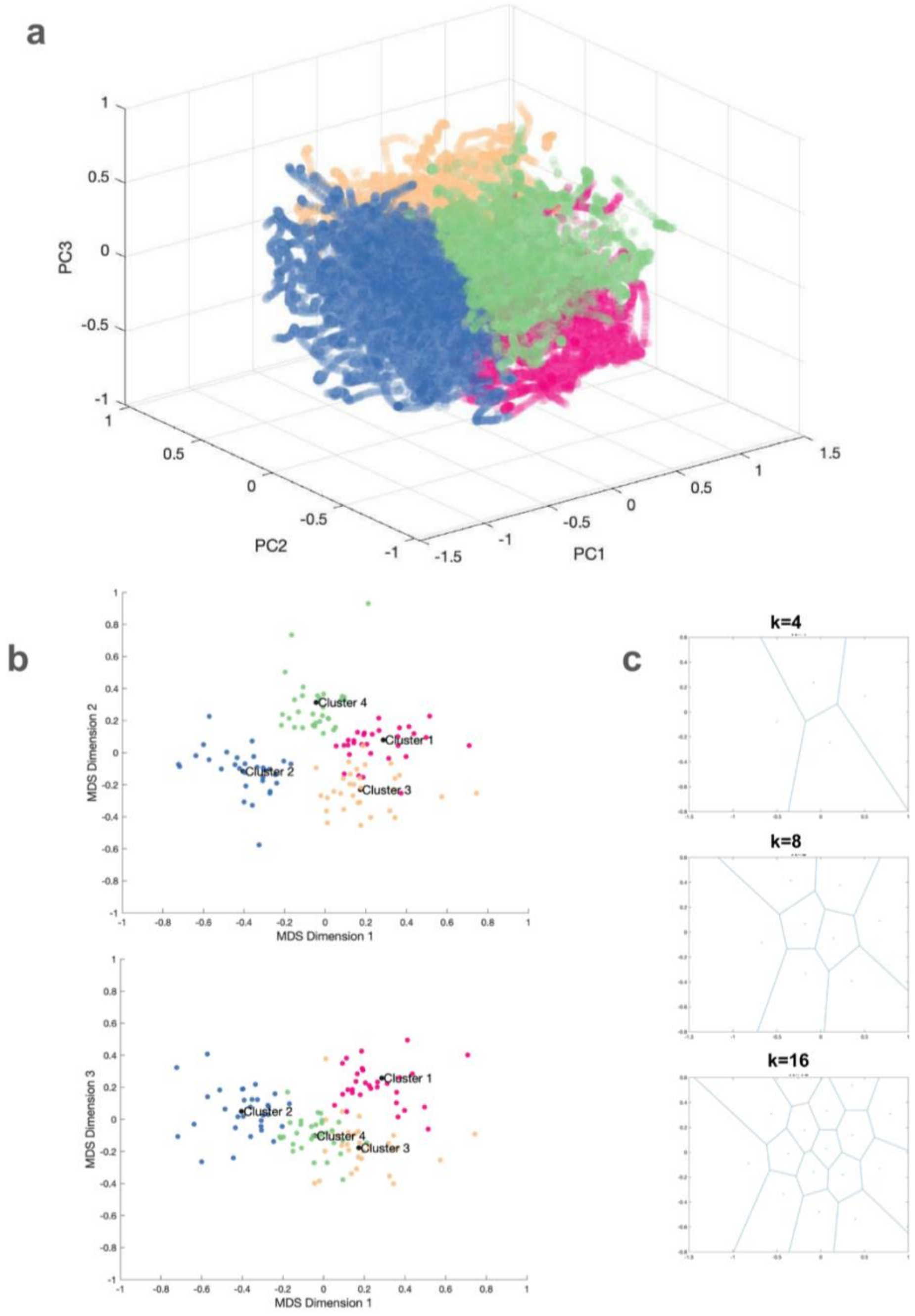


***Figure S3:*** *Partitioning of the twelve-dimensional time series data into k = 4 clusters.* ***a*** *The complete time series data from both meditation groups is shown. Each dot represents a twelve-dimensional time point (101 per session) in the first, second and third principal component space. Each dot is coloured according to the closest centroid in squared Euclidean distance.* ***b*** *for each participant, the values within each cluster are averaged to obtain a single 12-dimensional vector. Using Multidimensional Scaling the distances between these vectors (one per participant per cluster) are illustrated. The mean Euclidean distance between cluster 1 and cluster 2 is greatest (d=0.756), followed by the distance between clusters 2 and 3 (d=0.638). Cluster 1 is approximately equidistant to cluster 2 (d=0.5598) and cluster 3 (d=0.5569). Cluster 4 is approximately equidistant to cluster 2 (d=0.5949) and cluster 3 (d=0.595). Cluster 2 separates from the remaining clusters along the first MDS dimension, while clusters 4 and 3 separate along the second MDS dimension. Cluster 1 additionally separates from cluster 3 along the third MDS dimension.* ***c*** *Illustration of the partition of the same data in a 2D plane into k Voronoi cells, where each point in a given cell is closer to its centroid than to any other centroid.*
